# The neural computations underlying context dependent attribute-based valuation of complex stimuli

**DOI:** 10.1101/2025.10.10.681681

**Authors:** Aniek Fransen, Vincent Man, Kiyohito Iigaya, John P. O’Doherty

**Affiliations:** California Institute of Technology; Columbia University

## Abstract

Adaptive decision making requires value computations to be flexible because we often need to value a stimulus differently as the context changes. A stimulus might be highly desirable in one context but completely unappealing in another. However, it is not known how the brain can support such flexible modulation of overall stimulus value. Here we test a model of flexible value construction whereby individual attributes of a stimulus are converted into contextually dependent attribute-specific value representations before being combined into an overall integrated stimulus value. To test this framework, human participants (online n=95, MRI n=35) provided ratings for 75 unique high-dimensional clothing stimuli under three instructed ‘goal-contexts’ designed to elicit differences in overall value judgments for the items. Integrated value ratings for each stimulus were found to be goal-context dependent, while individual stimulus-attributes varied markedly in how they contributed to value ratings across goal contexts. In the fMRI data, representations of the absolute levels of particular attributes were revealed in visual areas. In contrast, encoding of individual attributes in value space, alongside integrated overall stimulus value, was present within distinct regions of prefrontal cortex. More specifically, behaviorally relevant attributes in value space and integrated stimulus value were found in vmPFC and dmPFC respectively. These findings indicate that the construction of value for high-dimensional stimuli is achieved through the computation of goal-context-dependent attributes in value space, providing mechanistic insight into how the brain can flexibly modulate stimulus-values as context changes.

## Introduction

Models of decision-making propose that the value of multi-attribute objects relies on the integration of their composite stimulus-intrinsic attributes (Kahnt et al. [2011], von Winterfeldt and Fisher [1975]). Most simplistically, the value of a gamble, for example, is computed by integrating the probability and magnitude of the prize on offer (Stewart [2011]). Similar principles have been proposed to facilitate value judgments for objects with high-dimensional attributes (Zysset et al. [2006]), where the presence and degree of various stimulus-intrinsic attributes are evaluated and subsequently summed in a weighted fashion to arrive at an overall object value (von Winterfeldt and Fisher [1975]). The resulting sum is then suggested to facilitate decision-making through between-stimulus comparisons. Previous research has examined the relationship between stimulus-intrinsic raw attributes and integrated value, demonstrating that a weighted combination of constituent attributes of a stimulus can predict how much value is assigned to a stimulus overall. For instance, the nutritive content of a food item can influence a person’s willingness to consume it (Suzuki et al. [2017]), while the visual properties of a piece of art contribute to aesthetic judgments (Iigaya et al. [2021]). This influence of composite attributes on value has been observed across various types of stimuli, ranging from t-shirts with aesthetic and semantic prints (Lim et al. [2013]) to gambles (Farashahi et al. [2019]).

However, focusing on only raw attributes and a fixed weighted integration of attributes to compute an integrated value has left an important gap in our understanding of the computations at play. This gap becomes clear when we consider that value judgments are not made in a vacuum but are highly flexible in that they can be heavily influenced by a person’s current goals (Castegnetti et al. [2021]), internal homeostatic state, or environmental context (Farashahi et al. [2019]). The same (visually identical) item can be valued highly under some circumstances yet considered low in value in other situations. For example, a ski jacket might be deemed highly valuable to an individual when going on a skiing holiday, but the same jacket would be of very low value when going on a beach vacation. A fundamental open question then is how can such highly flexible contextually-specific value computations be accommodated and understood within the framework of an attribute-based integration mechanism for computing overall stimulus value.

Here, we aim to address this question. We hypothesize that there exists a critical intermediate computational process between the representation of the attributes that make up a stimulus, and the overall integrated value. In this intermediate step, the value of individual attributes is computed in a context-sensitive manner. This context-sensitivity implies that the attributes of, say, a ski-jacket might be valued differently in different contexts. Our hypothesis proposes such context-sensitive attributes (e.g., its warmth) are evaluated in a context-dependent value space, before being integrated into an overall value signal. The goal of the present study is to investigate whether attribute representations in value space exist in the brain, as these would form an intermediate stage in the on-line computation of value for a given stimulus.

Much is known about the representation of individual stimulus attributes in the brain. In particular, distinct regions of prefrontal cortex have been implicated in different aspects of this process. Regions of the lateral prefrontal cortex, comprising both dorsolateral prefrontal cortex and lateral orbitofrontal cortex, have been found to encode information about individual stimulus attributes (e.g., Kahnt et al. [2011], Suzuki et al. [2017], Iigaya et al. [2023]). In contrast, the medial frontal cortex, particularly the ventromedial aspect, has been implicated in encoding the combined value of multi-attribute objects (e.g., Lim et al. [2013], McNamee et al. [2013], Iigaya et al. [2023]). However, in all of these studies it is not known how these brain regions can facilitate context-specific value computations through attribute integration. More specifically, these studies have not distinguished representation of raw stimulus attributes (i.e. the amount of attribute present in a stimulus irrespective of context), from the representations of individual attribute value. This gap means we do not yet know how multi-attribute value computation can be flexibly adjusted to changes in our goals, while this is a process that is abundantly present in our everyday decision making.

We designed a novel experimental paradigm specifically to address this question by distinguishing the neuronal representation of each of three distinct components: 1. the degree to which attributes are present in a stimulus (stimulus-intrinsic raw attributes), 2. the hypothesized value these individual attributes confer given the current goal (attributes in value space), and 3. the integrated value of the stimulus. We further aim to investigate the neural mechanism through which stimulus-intrinsic attributes and (if extant) attributes in value space contribute to the goal-dependent valuation of multi-attribute stimuli.

## Results

Here we set out to investigate the neural mechanisms underlying the computational steps that enable us to form goal-dependent values for multi-attribute stimuli. To achieve this, we designed a novel task; the “shopping task”. In this task, we introduced participants to instructed “goal-contexts”, enabling us to induce and measure changes in the flexible computation of goal-dependent values across goal-contexts. Under each goal-context, participants viewed 75 unique gender matched clothing stimuli. These goals included; going on a ski trip, visiting a tropical island and going for a job interview. Participants indicated how much they would like to wear the displayed item under the current goal (see Figure 1). Ratings ranged from 1-“do not want to wear at all” to 7-“want to wear a lot” (see Methods for more details). We expected that the subjective integrated value rating of a particular item (operationalized here by how much a participant would like to wear the item in a given goal-context), would be modulated by the goal-context (i.e., to be context-dependent). To reprise a previously introduced example, a heavy winter jacket might be highly valued in the ski trip goal-context (henceforth ‘context’), while being deemed of very low value in the tropical island vacation context. Similarly, a swimsuit might be highly valued in the tropical island vacation context, but of low value in the job interview context.

**Figure 1:**
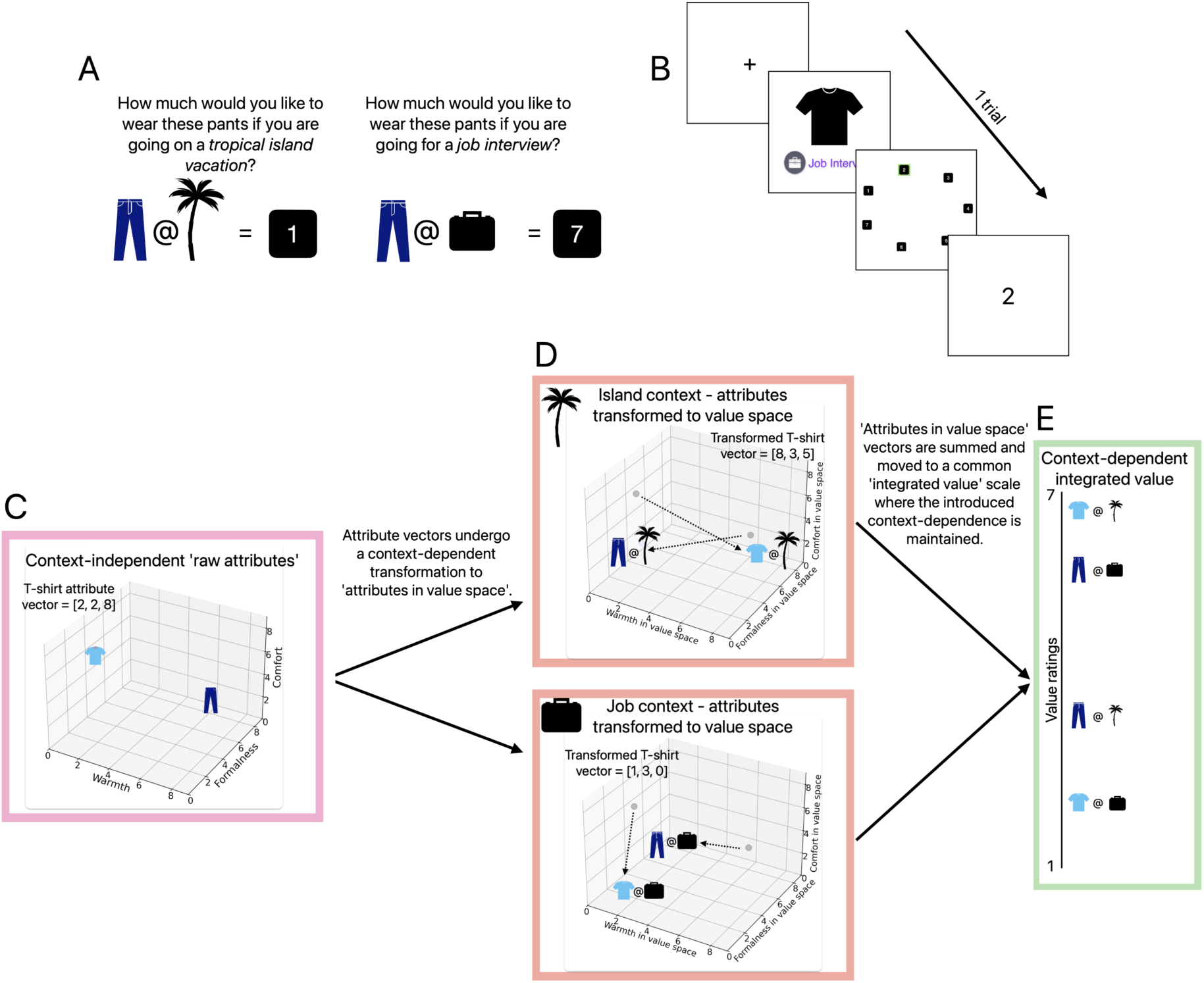
**A)** Participants performed a rating task under three instructed goal-contexts: tropical island vacation, ski trip and job interview. Participants indicate how much they would like to wear the clothing item shown given the goal on a scale from 1 through 7 (1; don’t want to wear at all, 7; want to wear a lot). We show participants the same item across all three goal-contexts to measure context-dependent shifts in valuation. **B)** One trial consists of four stages. Trials start with an inter-trial interval, followed by the item (t-shirt here) and the current goal-context. After observing the item and goal-contexts, participants are asked to indicate their rating. This rating is reported by navigating around a circular scale (current position indicated with a green highlight) in a clockwise or counter clockwise direction and confirming the rating with a button press. This manner of rating scale separates the magnitude of the rating from the number and direction of button presses. After confirming their rating, trials are concluded by showing participants the recorded rating (e.g., “2”). Image order was fully randomized and trials under a certain goal were presented in a blocked manner (15 trials before switching). Each fMRI run contained exactly one block of each goal. T-shirt shown is for illustration purposes only and was not part of the stimulus set (see Methods for details). **C)** Schematic representation of the three constructs under investigation: raw attributes (pink schematic), attributes in value space (red schematic) and integrated value (green schematic). We show two example stimuli (t-shirt and pants) and schematically represent the transformations these items undergo. Firstly, in raw attribute space (here shown in 3D: “warmth”x”formalness”x”comfort”) items location (and thus the raw attribute axes) are context-independent. Item location indicates the presence of a certain attribute in the clothing item. **D)** The raw attribute vectors then undergo a context-dependent transformation to ‘attributes in value space’. ‘Stimulus @ context’ indicates the context-dependence of these new axes. Arrows are shown to schematically indicate this transformation from raw attributes to attributes in value space. Item location in 3D-attribute-in-value-space indicates not only the amount of an attribute that is present in the clothing item but also the current goal-dependent value of this attribute. As such attribute in value space vectors are different across different contexts (here shown island and job contexts). **E)** Lastly, item’s attributes-in-value-space vectors are summed together and moved to a common rating-scale. We schematically show this subjective integrated value axis with each of the example items under each goal-context. The integrated value is directly measured in the task through the participant stimulus ratings (see **A** - on a scale from 1 through 7). Note here that the same item shown under different goal-contexts occupy distinct locations along the integrated value axis to indicate the goal-context dependency of the integrated value ratings.

We leverage this new paradigm in two experiments: one collected in an online sample (Prolific; n=95, 57 female) and one conducted in an fMRI scanner (n=35, 20 female, age 18-64). As part of both of these experiments, we measured subjective stimulus ratings directly through the described rating task (see Figure 1).

Using this task, we firstly aimed to capture the neural representation of the raw attributes that make up a stimulus. These raw attributes quantify the perceived or objective content of a given attribute in the stimulus as a whole (independently of that attribute’s value). For instance, a heavy jacket will be ‘high’ in the attribute of warmth, regardless of whether it is worn on a ski trip or on a tropical island vacation (see Figure 1C). In addition to the neural computations underlying raw attributes, we set out to find neural representations of attributes transformed into value-space. These ‘attributes in value space’ can be distinguished from raw attributes, since the warmth attribute will be valued differently depending on the context (e.g., whether an individual is wearing the jacket on a ski-trip or on a tropical island vacation - see Figure 1D). Finally, our task allows us to also probe the subjective (integrated) value of such multi-attribute stimuli (see Figure 1E). The integrated value is the value of a stimulus (e.g., the heavy jacket) within each context, when taking into account multiple attributes simultaneously.

Given our main objective is testing for evidence of the encoding of attributes in value space, we picked a small set of salient high-level attributes pertinent to value decisions about clothing items: the “warmth”, “formalness”, “comfort” and “festiveness” of the items. While these are a limited subset of a much larger set of possible attributes, they are sufficient to allow us to distinguish raw attribute, attributes in value space and integrated value representations. We also examined “low-level”visual attributes of the clothing items which correspond to elementary visual properties of the stimuli (see Methods for more details and Iigaya et al. [2020]). We confirm our claims of sufficiency through this set of “low-level” attributes and contrast the neural representation of these lower level attributes with the selected higher level attributes.

### Behavioral contribution of attributes to integrated value ratings

We started by probing the relationship between attributes and value at a behavioral level. As attributes in value space (see Figure 1D) are latent variables, we sought to approximate their effect on integrated value through a weighting of raw attributes. This weighting can be thought of as an estimation of the transformation the raw attributes undergo to hold value-based information (see Figure 2C to D).

**Figure 2:**
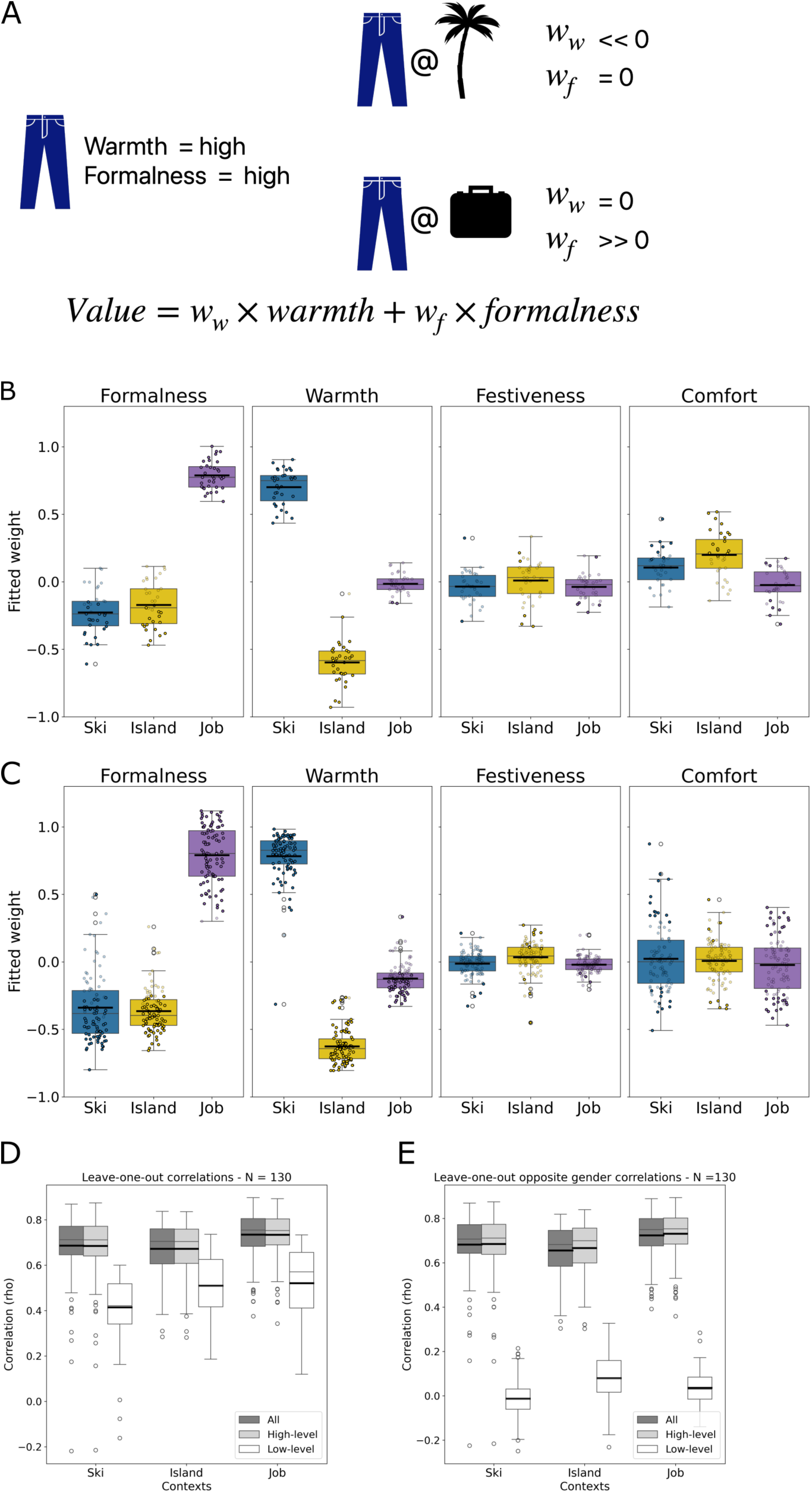
**A)** Schematic representation of the data used for the behavioral model fitting procedure. Participant ratings (i.e., subjective integrated value) for the same item will vary across goal-contexts while the raw attributes are context-independent. We expect differences in the context-specific weights (*w_warmth_, w_formalness_, .., w_n_*) to capture the differences in individual raw attribute contributions to the overall within context stimulus ratings. As such, we are plotting these raw attribute weights, fitted in each of the three contexts, below. **B)** Attributes’ influence on fMRI participant’s (n=35) ratings varied greatly depending on the goal. For each participant we fit a GLM to predict participants ratings under each goal-condition. We included each of our apriori raw attributes as regressors to probe their goal-specific contribution to the overall integrated value rating reported by the participant. We are plotting the extracted beta-coefficients for each of these raw attribute-regressors. The resulting distributions of beta-coefficients across subjects are shown in box and whisker plots. For each box and whisker plot, the middle line denotes the median, and the bottom and top edges of the box indicate the first quartile (Q1) and third quartile (Q3) of the distribution. The ends of the whiskers represent the maximum and minimum data points not considering outliers. Outliers are shown in open circles and are denoted as such when they surpass either the lower bound (*Q*1*—* 1.5*\**(*Q*3*—Q*1)) or the upper bound (*Q*3+1.5*\**(*Q*3*—Q*1)). Across subject means are reflected in bold horizontal lines. Additionally, individual subject model coefficients are plotted with the significance of the coefficient indicated through its color fill (white for non-significant and black for significant regressor coefficients; threshold used *p <* 0.05). **C)** Results for the identical analyses conducted on the online sample (n=95). We observed a highly similar pattern in raw attribute contribution across subjects and goal-contexts. Box and whisker plots mirror those described above and individual regressor coefficients are similarly shown as scatter (with fill indicating significance as above). **D)** Model evaluation results for raw attribute-based value prediction. Box and whisker plots reflect the spearman correlation between the model predicted value rating and subjective ratings of a left-out subject. We trained an elastic net regression model to predict subjective value ratings based on varying sets of raw attributes. Darkest boxes reflect model performance for models including all raw attributes. Middle boxes indicate the performance of models trained on the a priori raw attributes, while rightmost light colored boxes show the performance of models exclusively trained on computer vision extracted low-level attributes. Models are trained across all (both samples) but one left out subject, who’s ratings are subsequently predicted by the model. This procedure is repeated 130 times for each model, such that each subject is left-out once. Model performance is statistically identical across models including all and high-level a prior attributes (Mann-Whitney U test: ski; stat=8387, *p* = 0.92, island; stat=8457, *p* = 0.99, job; stat=8424, *p* = 0.97. Model performance is however statistically different for models including only low-level computer vision extracted attributes (Mann-Whitney U test: ski; stat=1595, *p* = 3.58*e^−^*^35^, island; stat=13896, *p* = 2.66*e^−^*^19^, job; stat=15262, *p* = 2.75*e^−^*^29^). **E)** Similarly to above we evaluate model performance across models trained on various sets of attributes. Here however, the data is split across genders. The model is trained on one gender, while the left-out subject is from the opposite gender. Importantly, subjects with different genders observe two unique sets of stimuli allowing us to evaluate our models on a completely left-out stimulus set. Spearman correlations are computed for the model predicted value rating versus subjective value ratings for each left-out opposite gender subject. Equivalently to D, each box and whisker plot includes 130 models (each subject is left out once). Across-gender models trained on all raw attributes and those trained on high-level a prior raw attributes perform equally well (Mann-Whitney U test: ski; stat=8616, *p* = 0.78, island; stat=8995, *p* = 0.37, job; stat=8875, *p* = 0.48). We again do observe a significant difference in performance between models trained on high-level raw attributes and those trained on only low-level raw attributes (Mann-Whitney U test: ski; stat=16764, *p* = 8.57*e^−^*^43^, island; stat=16896, *p* = 4.16*e^−^*^44^, job; stat=16900, *p* = 3.80*e^−^*^44^).

We expect our a priori selected attributes to be a statistically significant contributors toward the overall value in at least one context. If met, this expectation verifies that the attribute indeed contributes to the overall value of a stimulus, in a manner consistent across individuals. To test this, we evaluated the independent main effect of each attribute under each context through a mixed-effects modeling approach that controls for individual differences in stimulus ratings (see Methods for mixed-effects model specification - fMRI sample only). Confirming the relevance of our chosen attributes to the computation of participant ratings (i.e., integrated value), we found that all attributes are significant independent predictors of value under each of our instructed contexts (T-tests: see Supplemental Table S1 for attribute-context statistics).

Secondly, we would expect to observe differential transformations of raw attributes as they turn into attributes in value space and contribute to the goal-dependent integrated value (i.e., stimulus ratings). To test if such a goal-dependent transformation indeed exists, we implemented context and participant specific regression models to extract weights for each of our a priori raw attributes (see Methods for details - and supplemental Figure S2G). We expected attributes to contribute differently to stimulus ratings depending on the goal-context. To test this, we directly evaluated if the coefficients of attribute main effects are statistically different between contexts. If we find across-goal differences between fitted coefficients, this would provide evidence that the estimated attribute contributions are indeed goal-dependent. We found this to be consistently true for formalness (Kruskal-Wallis; fMRI: H=70, *p* = 6.23*e^−^*^16^, Online: H=189, *p* = 1.12*e^−^*^41^) and warmth (Kruskal-Wallis; fMRI: H=92, *p* = 1.01*e^−^*^20^, Online: H=249, *p* = 8.74*e^−^*^55^) but not for festiveness (Kruskal-Wallis; fMRI: H=5, *p* = 0.07, Online: H=28, *p* = 7.08*e^−^*^7^) and comfort (Kruskal-Wallis; fMRI: H=32, *p* = 9.02*e^−^*^8^, Online: H=1, *p* = 0.51). This result highlights that while goal-dependent attribute influence exists, this is not necessarily true for every attribute in every participant. Note that in the online study (see Figure 2C) we did not obtain subjective attribute ratings. In this sample we therefore instead used the stimulus-specific attribute means across fMRI subjects for this analysis. We additionally replicated these results with weights fitted using a logistic regression model (see supplemental Figure S2G), with the only qualitative difference being a significant effect now also emerging for comfort in the online group (*p* = 0.03, see supplemental Table S2 for all statistics).

### Integrated value predictions based on high-level, but not low-level, attributes generalize to left-out stimulus sets

Next, we aimed to evaluate the degree to which our a priori (high-level) raw attributes can predict value ratings across individuals and stimuli. If our set of attributes has a generalizable effect on integrated value we can feel confident we have indeed selected a sufficient set for our following analyses. Additionally, across-subject generalization would let us conclude there is a level of communality to the computations underlying valuation across subjects in our sample, which will be helpful in analyzing the neural data specifically. Previous research has shown that low-level visual attributes can also contribute to valuation of multi-attribute stimuli. We therefore also extracted a set of data-driven low-level attributes using computer-vision methods following a previously validated approach (see Methods and Iigaya et al. [2021]). For comparison we tested models with 1) the low-level attributes alone, 2) the high-level attributes alone, and 3) both low- and high-level attributes. We hypothesized that high-level attributes, by virtue of their more abstract nature, possess a better capacity for generalization across individuals compared to low-level attributes. To investigate the distinct contributions of high- and low-level raw attributes, we employed a cross-validation (leave-one-subject-out) elastic net model across both experiments (see supplement Figure S3A and B for separate results). Across held-out participants, we see that a model including the high-level raw attributes alone, predicts equally well as a model including all raw attributes (i.e., both high and low level) (spearman correlation: mean rho=0.698 and rho=0.697 for all and just including high-level attributes respectively; Mann-Whitney U test: stat= 8387, *p* = 0.92 for ski; stat = 8456.5, *p* = 0.99 for island; stat= 8424, *p* = 0.97 for job). This stands in contrast to models trained only on the low-level raw attributes, which yield a significantly worse performance than those trained on high-level raw attributes (Mann-Whitney U test: stat=1595, *p* = 3.58*e^−^*^35^ for ski; stat=13896, *p* = 2.66*e^−^*^19^ for island; stat=15262, *p* = 2.75*e^−^*^29^, see Figure 2D). As such we conclude that the high-level attributes are better able to predict value than low-level attributes and do so in a manner that captures similarities across participants.

Next, we aimed to further investigate how well a model trained to predict value ratings, based on either high or low-level attributes, can generalize such predictions. Here we test these models’ value predictions for both an unseen stimulus set, and in a completely separate group of individuals. We leveraged the fact that male and female identifying participants were presented with two distinct stimulus sets with male and female gendered clothing items respectively. We exploited this design feature by training our model on one gender and testing it on the ratings from individuals in the opposite gender group, who thus saw the other set of stimuli. Comparing the generalizability of high-level and low-level attributes through this approach shows a stark difference. While a model trained on the high-level raw stimulus-attributes generalized well across stimulus sets (and thus genders), a model trained only on the low-level raw attributes failed to generalize (see supplement Figure S3D and E for separate online and fMRI subject results). This difference was robustly significant (Mann-Whitney U test: ski; stat=16764, *p* = 8.57*e^−^*^43^, island; stat=16896, *p* = 4.16*e^−^*^44^, job; stat=16900, *p* = 3.80*e^−^*^44^, see Figure 2E). These findings indicate a key advantage of high-level attributes and highlights their strength in predicting the value of stimuli. By virtue of the abstraction of these attributes from the basic visual properties of a stimulus, they are able to capture relationships between attributes and value that generalize well across stimulus sets and groups of individuals. To gain insight into the neural representation of these raw attributes, we next turned to the fMRI data.

### Visual cortex represents low- and high-level stimulus-intrinsic attributes

We expect the brain to separately represent each of the three constructs under investigation (raw attributes divided into low and high-level, attributes in value space, and integrated value ratings), as they make up distinct component processes underlying flexible valuation of multi-dimensional stimuli. We first tested the encoding models of raw attributes. We hypothesized that, even though high-level attributes are sufficient to predict valuation, the brain represents both low-level and high-level attributes. Additionally, we expect such representations to be context-independent since the raw attributes are static properties of the stimuli.

The whole brain analysis shows that these raw attribute representations are concentrated in the visual cortices (F-test across high-level attributes shown in red in Figure 3B - with max voxel for high-level raw attributes: [56,15,32], cluster corrected FWE*<*0.05, voxel threshold *p <* 0.001 for all reported clusters). Consistent with previous findings (e.g., Iigaya et al. [2023]), early visual regions tend to encode low-level attributes more prominently than high-level attributes. In contrast, we observe that higher visual regions more strongly represent high-level visual attributes (see Figure 3C and Figure S5).

**Figure 3:**
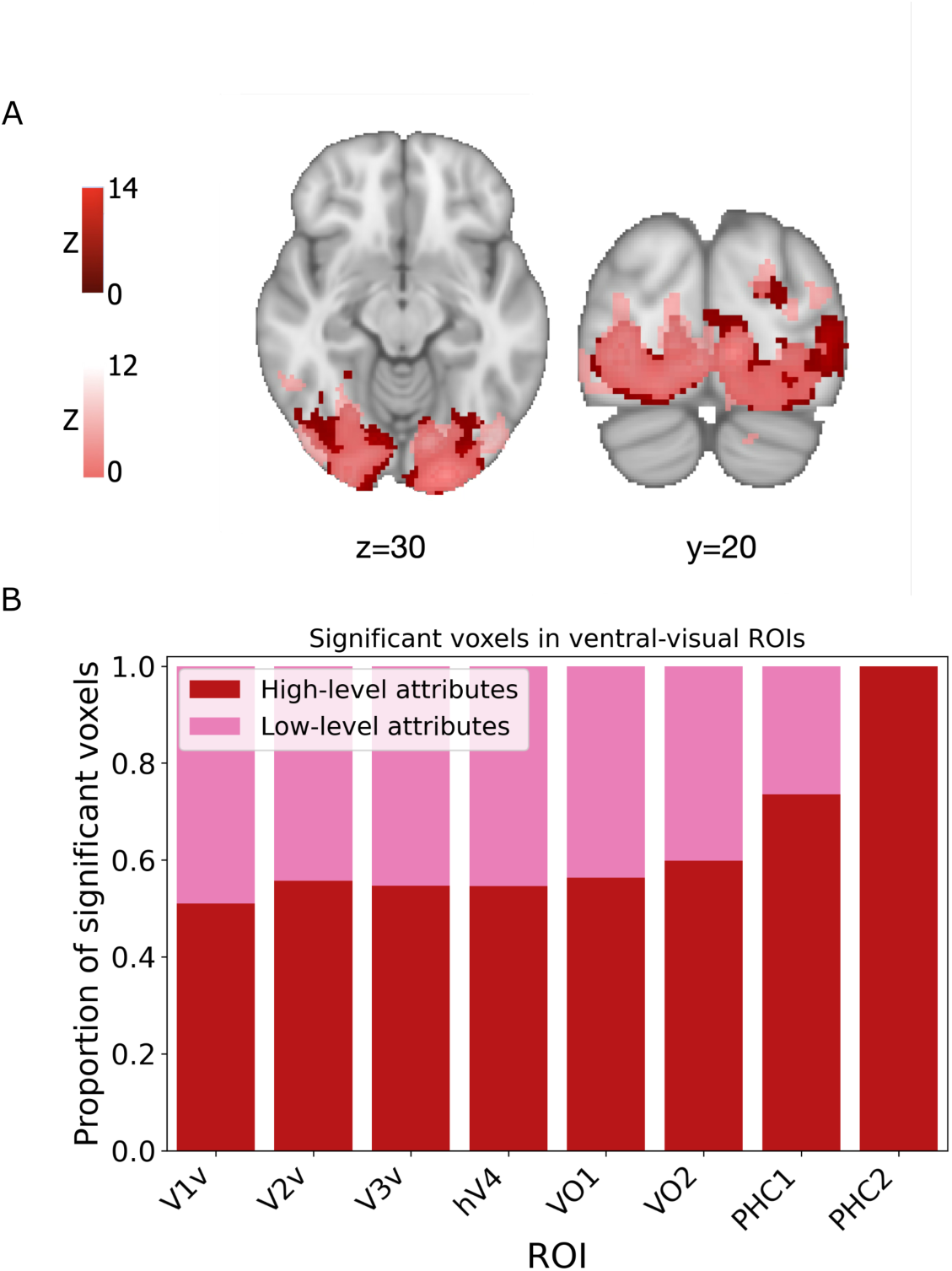
**A)** Since raw attributes are context-independent we can model their presence (e.g., the amount of warmth provided) in one model across contexts. In practice, we test for such context-independent raw attribute representations through fitting an encoding model. We observe that both computer-vision extracted attributes (F-test across attributes shown in pink) and participant-rated attributes (F-test across attributes shown in red) are represented in overlapping parts of the occipital cortex (cluster corrected FWE*<*0.05, voxel surpassing *p <* 0.001). **B)** When plotting the proportion of significant voxels (before cluster correction) that represent low or high-level attributes across ventral-visual cortex ROIs we observe a gradually more prominent representation of high-level attributes along the posterior-anterior axis.

### Frontal cortex is a candidate region to hold attributes in value space signals

We next aimed to test the hypothesis that the brain represents attributes in value space (see Figure 1D), in addition to the stimulus-intrinsic raw attributes shown to be encoded above. If a brain area encodes raw stimulus attributes then the representation of that attribute would not be expected to change as the goal context changes. In other words, the raw attribute would be represented in the same way across all goals as it should be goal-context independent (see Figure 4A). Alternatively, if an attribute is being represented in value space then it would be expected to exhibit differences in its representation as goal-context changes. Such context-based representational shifts are expected especially in the case where the attribute is relevant for the goal-value computation (such as in the case of the warmth attribute when computing the value of a ski-jacket vs a swimsuit) (see Figure 4A). To distinguish between these possible coding schemes, we set up an attribute decoding generalization analysis. We first trained a regularized classifier to decode each of our subjectively rated raw attributes under a specific goal. For example, we might train a model to classify whether clothing stimuli shown under the ski goal are high or low in warmth (median split, for more details see methods). We can then test this model on the same goal (ski) or on one of our two other goals (job interview or island vacation).

**Figure 4:**
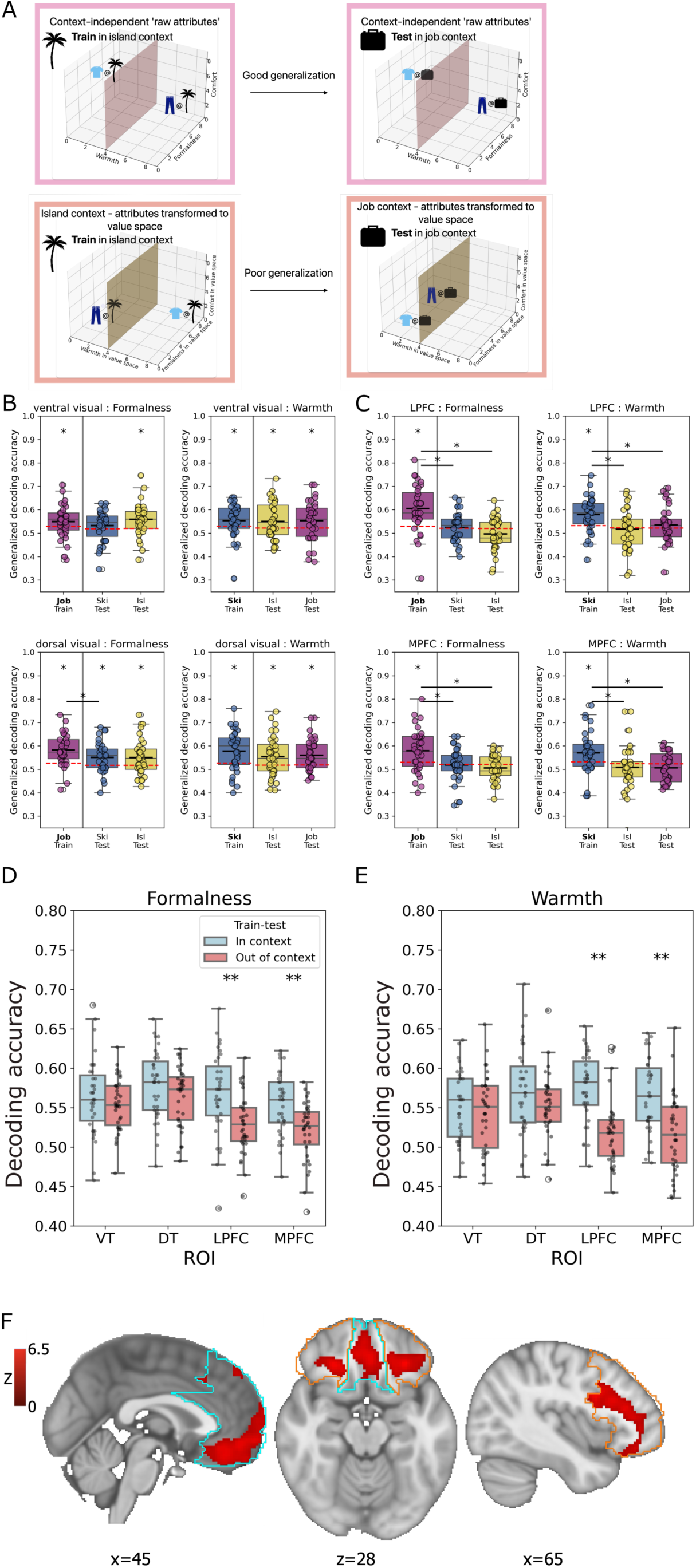
**A)** Schematic representation of model performance interpretation. If context-independent raw attributes are represented in an ROI we would expect good generalization performance. Since raw attribute representations are context-independent the hyperplane fitted in one context (e.g., separating high and low warmth items under island) will similarly separate raw attributes under another goal-context (i.e., job). However, if instead an ROI represents context-dependent attributes in value space we expect poor generalization performance. As shown in the example schematic, the value axes shift between contexts and the items shift position accordingly. Therefore a hyperplane fit to separate items in one context (e.g., island) will be unable to separate those items under a different context (e.g., job). **B)** Within subject ventral and dorsal visual cortex generalization decoding accuracies for formalness and warmth attributes. The goal-context used for training is shown in the left most position (as annotated on the x-axis) and dashed lines show permuted chance level (mean of null) (see Methods). Statistical significance of the context-specific model performance is computed with respect to each context-specific null and any significant results are indicated (*\**: *p<* 0.05). We additionally compute statistics for the pairwise comparison between the training and each testing context. The significance of these between-context decoding accuracy differences are similarly indicated (*\**: *p <* 0.05). For both ventral and dorsal visual cortex ROIs we observe significant within and out-of-context model performance in addition to no significant differences in decoding accuracies between training and texting goal-contexts for both formalness and warmth attributes. **C)** Similarly, we show the generalization decoding for lateral and medial prefrontal cortex (lPFC, mPFC) ROIs. Here we do observe model performance is not different from chance in the out-of-context test sets. Additionally, we do observe significant differences in decoding accuracies between training and testing goal-contexts. This implies a shift in the underlying neuronal representation of the attribute across goal-contexts. **D)** Formalness decoding accuracy computed for each of our ROIs (VT: ventral temporal, DT: dorsal temporal, LPFC: lateral prefrontal cortex, MPFC: medial prefrontal cortex). Decoding accuracy of out-of-context models is shown in red bars with within-context model performance shown in blue for comparison. Importantly, this figure includes model performances across all trained and tested models. Within-context models thus include those for the ski, island and job contexts with each of these models evaluated across the other two contexts. Individual dots shows within subject average performance across the three within-context models (under blue bars) and the within subject averaged out-of-context model performance (under red bars). T-tests are used to compare this out-of-context performance with the within-context performance for each ROI (*\*\**: *p <* 0.01). **E)** Similarly for the warmth models. We again observe significant differences in average model performance within contexts and out-of-context in frontal areas (LPFC and MPFC). **F)** The average encoding across estimates of attributes in value space is shown to be significant in various areas within our lPFC (outlined in orange) and mPFC (outlined in light blue). These clusters are focussed on areas including the vmPFC stretching into medial OFC, lateral OFC and more dorsal areas of lPFC. All result are cluster corrected within the ROIs for FWE*<*0.05 with voxels surpassing *p<* 0.001.

If we find that a model trained on one goal’s raw attribute ratings generalizes well to another goal this means the underlying neural activity pattern across goals is consistent, as expected when representing a raw attribute. For example, the brain signal encoding the raw attribute of high warmth would be similar for both the ski trip and island vacation goal-contexts. Such similarity in the neural representation of a raw attributes across goals would exclude the possibility of the attribute being represented in value space. Alternatively, if a decoder successfully decodes a raw attribute in the training context, yet performs poorly when tested on another goal context, that would indicate that the neural representation of the raw attribute has shifted from one goal to the next (see Figure 4A). Such a decrease in decoding accuracy would be expected if the area computes attributes in value space, as this is a goal-dependent signal. Put more concretely, if the model trained on warmth under the ski goal has poor performance when tested on the tropical island goal this would be indicative of distinct neural coding axes for high vs low warmth across these goals. Evidence of distinctive neural representations across goal contexts would be suggestive of the type of goal-context-dependent modulation of neural activity necessary for the brain representation of attributes in value space. We can then leverage this evidence to further investigate value-dependent neural codes.

Probing the generalization performance of the two attributes that displayed the most homogeneous behavioral pattern across subjects (formalness and warmth, see Figure 2), we observe consistent significant decoding both within the training and each of the test-contexts for the two visual ROIs (see Figure 4B). However, for the frontal ROIs (LPFC and MPFC) we observe significant differences in accuracy between the training and test contexts (see Figure 4C). Additionally, we find that in all test-contexts, this decrease in performance is so drastic the raw attribute can no longer be significantly decoded in the test context. These frontal ROIs thus display a shift in the representation of raw attributes from one goal to another, thereby meeting the criteria we identified as being necessary for brain areas to represent attributes in value space (see Supplemental Figure S6 for other attributes). We additionally probe the generalization performance across all train-test context combinations. We find the out- of-context decoding accuracy (see Figure 4D&E - in red) is significantly lower than the within context decoding accuracy in lateral and medial prefrontal cortex (see Figure 4D&E -Wilcox test; formalness: LPFC: *p* = 0.0009, MPFC: *p* = 0.0005; warmth: LPFC: *p* = 2.27*e^−^*^6^, MPFC: *p* = 2.87*e^−^*^5^). We ran these same generalization analyses for ventral and dorsal visual stream ROIs. Given our previous encoding analysis finding these regions represent raw attributes (see Figure 3A), we expect to find good generalization in these ROIs. We indeed observe no differences in decoding performance for raw attributes across the ventral visual-temporal (VT) and dorsal visual-temporal (DT) ROIs (Wilcox test; formalness: VT: p=0.20, DT: p=0.10; warmth: VT: p=0.29, DT: p=0.10).

### Prefrontal cortex represents attributes in value space through encoding of model estimates

While the generalization analysis confirms context-dependent attribute representations exist, this does not yet prove that this is a value-related context dependence. We therefore turn to our earlier behavioral models which computed the relationship between raw attributes and participants’ stimulus ratings across goal conditions. From these models we obtained estimates (i.e., *ω* weights) of the transformations necessary to compute attributes in value space - i.e., the transformation of raw attributes into value space. We now directly apply these estimated transformations to compute estimates of attributes in value space. To achieve this we multiplied subject and goal specific behavioral GLM *ω*-weights by the subjective raw attribute ratings to compute estimates of subjective attributes in value space (i.e., *w_w_ × warmth*). Attributes in value space should capture both the presence of an attribute in an item and the current value of that attribute. For example, a sweater will provide a lot of warmth (e.g., subjective raw warmth rating of 8) and when presented with this sweater under the goal-context of a ski trip, this warmth will be a large positive contributor to the overall item value (e.g., subjective *ω* of 0.6, leading to an attribute value of 8 *×* 0.6 = 4.8). While for the same sweater under the island vacation goal-context a negative attribute value would be estimated (e.g., subjective *ω* of -0.5: 8 *× \**0.5 = *\**4). These multiplications of fitted weights with raw attributes, thus leaves us with direct personalized estimates of attributes in value space. Next we will use these estimates to test for the encoding of attributes in value space using our neural data (see Methods for more details).

We restrict our analysis here to the areas shown to be candidates for representing attributes in value space in the generalization analysis above (LPFC and MPFC). When mapping out the clusters that correlate with our attribute- in-value-space estimates, we found significant clusters in both lateral and medial PFC (mean across attributes, see Figure 4F). When zooming into these large ROIs to delineate effects in individual sub-regions of MPFC and LPFC, we found significant encoding of attributes in value space in multiple PFC subregions including the medial and lateral orbitofrontal cortex (OFC) and parts of the ventromedial prefrontal cortex above the orbital surface (vmPFC) (max voxel in LPFC: [24, 80, 42], z=6.4). The abundant encoding of attributes in value space confirms that attributes in value space computations are indeed part of the neural mechanism underlying flexible value computation for multi-attribute stimuli.

### Subjective value ratings are represented across-contexts in widespread prefrontal cortex ROIs

We next turn to investigate the neural representation of the last construct under investigation; subjective integrated value ratings. First, we use the context-dependent stimulus value ratings reported by participants to train a decoding model (see Methods). We found that we could significantly decode participant ratings (i.e., integrated value) across all prefrontal sub-ROIs (see Figure5C - and see supplemental Figure S7B&C for visual ROI decoding results). Additionally, we probed the generalizability of the stimulus value ratings signal. To facilitate decision making (i.e., to make a comparison between options), we expected that the neural representation of such value ratings should generalize well across goal-contexts, as it has been abstracted away from the specific stimulus and goal under consideration to a common axis for valuation. We indeed find good generalization of the integrated stimulus value signal across contexts in frontal ROIs (see Figure 5B - and see supplemental Figure S7A for visual ROIs). Importantly, the ability of these frontal ROIs to represent a generalizable value rating signal also confirms the goal-specific raw attribute results discussed above do not stem from an inherent property of the ROI to switch representations across contexts.

**Figure 5:**
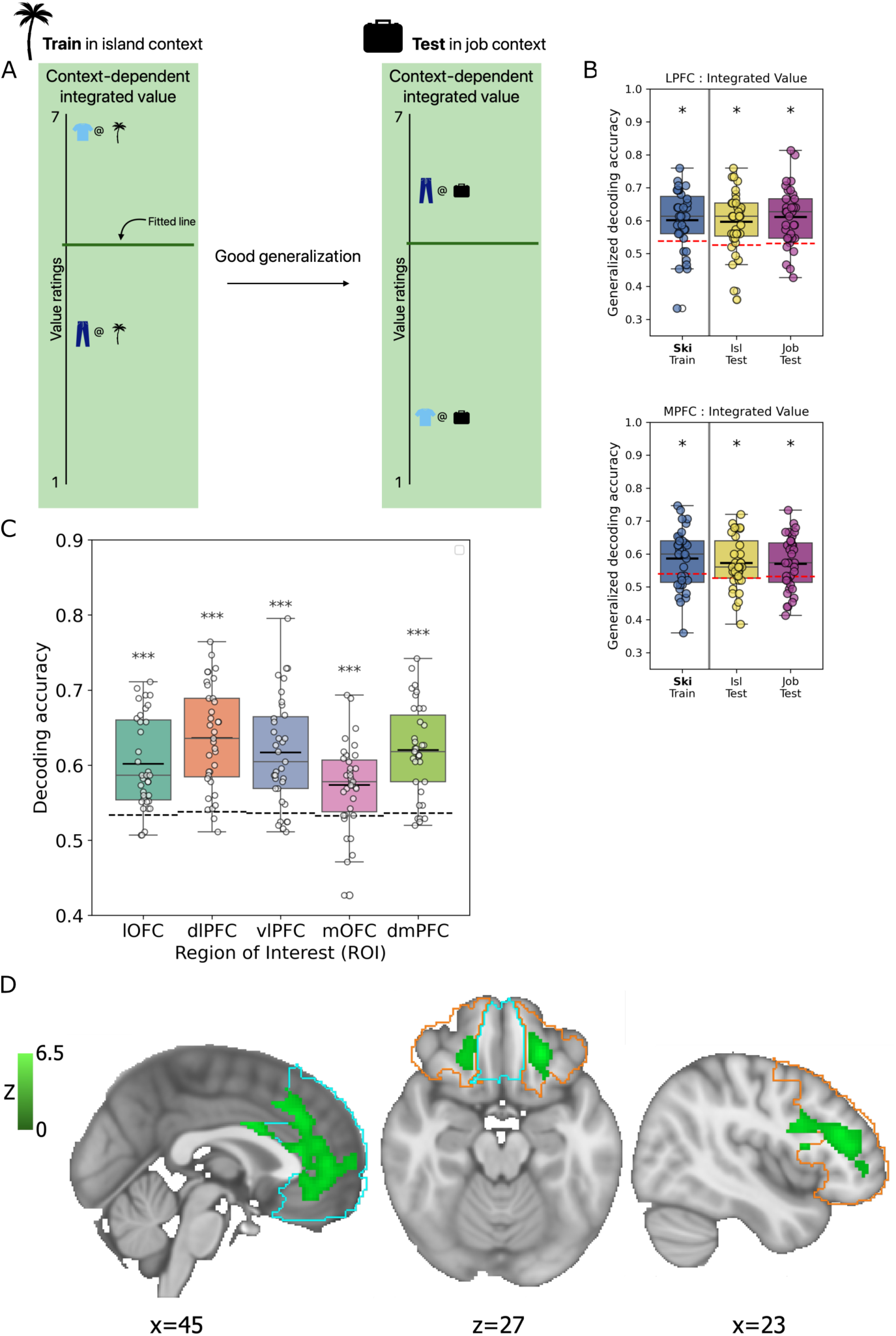
**A)** As items’ attributes in value space have been summed together to compute integrated values we once again expect good cross-context generalization. Importantly, at the level of integrated value identical items will have different ratings across contexts we expect items with a similar rating to be similarly represented in the brain. As such, at this level of computation, individual attributes of the clothing items are no longer represented and instead a common value scale allows for comparison across items and contexts. **B)** We indeed find good generalization of participants’ stimulus ratings (i.e., integrated value) across areas of prefrontal cortex. There are no significant between context differences in decoding accuracies and models trained on context show significant model performance on the two left-out contexts. Models trained on ski-context are shown as exemplars here (see supplemental figure x for other training contexts). **C)** In addition to a generalization decoding analysis, we performed integrated value (operationalized as stimulus ratings) decoding across the whole task. This decoding analysis was performed across sub-ROIs within the prefrontal cortex. We find universal significant representation of integrated value across frontal-cortex ROIs (*\*\*\**: *p<* 0.001). **D)** An encoding analysis confirms stimulus ratings are significantly represented in clusters that span areas of dorsomedial prefrontal cortex and ventral-lateral prefrontal cortex. We additionally find participants’ stimulus ratings are encoded in lateral OFC. The encoding model was ran within LPFC (outlined in orange) and MPFC (outlined in light blue) ROIs and result are cluster corrected within the ROIs for FWE*<*0.05 with voxels surpassing *p<* 0.001.

Similarly to the approach for attributes in value space, we next turned to model the integrated value (operationalized as participant stimulus ratings) in an encoding analysis performed in the LPFC and MPFC. We found participant ratings correlate with the neural signal along the medial wall of the prefrontal cortex stretching towards the anterior cingulate cortex (max voxel in LPFC:[56,81,28], z=6.5 - see Figure 5D). In the lateral prefrontal cortex, we find the strongest representation of participant value ratings in posterior parts of the ROI. Lastly, we find significant value rating encoding in lateral orbitofrontal cortex (OFC).

### Overlapping and separable encoding of subjective value ratings and attributes in value space in prefrontal cortex

We have found evidence that the brain represents both attributes in value space and subjective value ratings. Here we turn to address the open question of how the computation of these two value signals interact. We defined integrated value as the linear sum over attributes in value space (see Figure 1E and 2A and in line with previous research e.g. Iigaya et al. [2021]). Given this close relationship between these two constructs (one being the sum of the others) we might predict that the neural representation for these two constructs is similar as well. However, if we cannot find separate neural correlates for these two value computations we might not feel confident they are indeed two separate steps in hierarchical multi-attribute value computation. We therefore tested the extent of the similarity between integrated value ratings and attributes in value space. We employ a voxel-wise model in which the subjective stimulus ratings regressor directly competes with attributes in value space regressors to explain variance in the BOLD signal. In the most extreme case where attributes in value space are not computed independently from integrated value, we would expect to see the same voxels represent both variables. Alternatively, if we observe significant effects of attributes in value space or subjective integrated value ratings in this model these results would reflect each term’s unique variance (see Methods for more details). Such independent neural representation would imply that there is variance unique to the subjective stimulus ratings provided by our participants that cannot be captured by the equal linear combination of the four included attributes in value space. Using this combined model, we find substantial significant clusters of both subjective stimulus ratings and attributes in value space. Additionally, these clusters appear to be located in areas similar to those found in the models that focused on each construct separately (see Figure 6). We observed a similar ventral-dorsal separation in the prefrontal cortex ROI along the medial wall, with ventral correlates to attributes in value space. This ventral-dorsal pattern appears to be reversed within the lateral prefrontal cortex, in that the stimulus rating cluster occupies the posterior ventral LPFC. One major difference to note here is that in the combined analysis we no longer find significant stimulus rating encoding in lateral OFC but solely find evidence for attributes in value space across OFC regions. This suggests the lateral OFC representation of stimulus value rating (see Figure 5D) is now better explained by the attributes in value space regressors. This finding hints at the high similarity between the neural signal for integrated value ratings and attributes in value space in the region.

**Figure 6:**
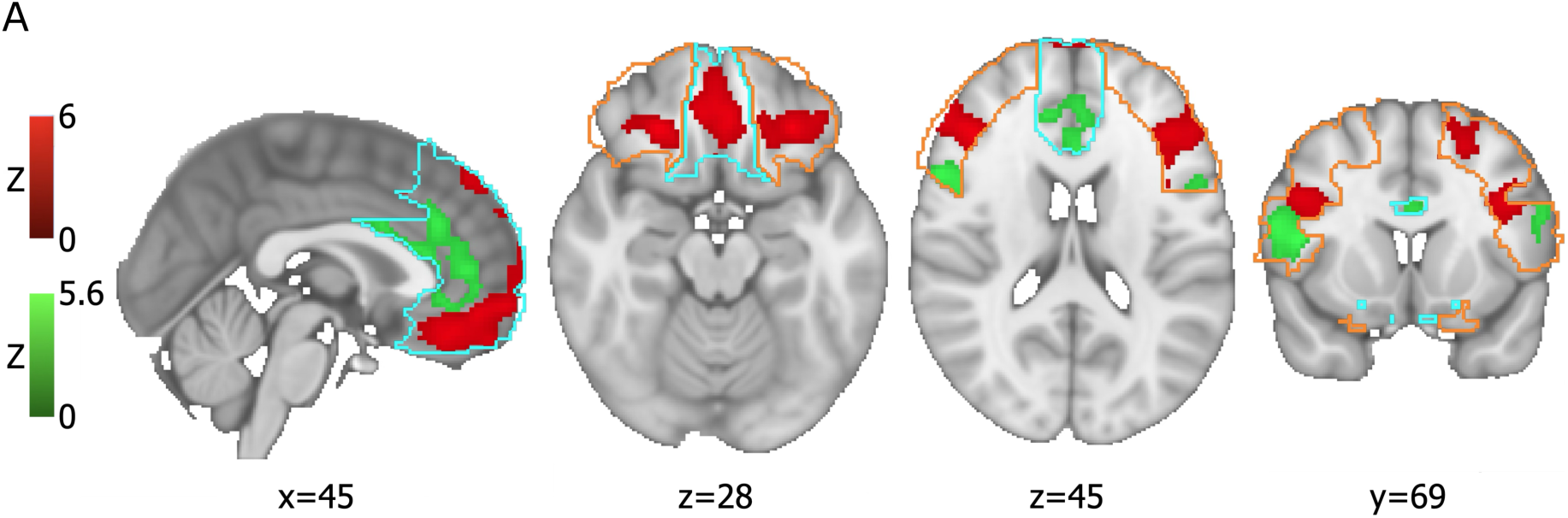
**A)** Encoding results of a model including both integrated value and attributes in value space. Since these results are from one unified encoding model they can be directly compared. Additionally, only variance unique to each of these two constructs is modeled here. Interestingly there are a lot of clusters that remain similar compared to Figure 4F and Figure 5D hinting at substantial unique variance in the neural representation of these constructs. However, we also observe differences, with most noteworthy a lack of integrated value encoding in the lateral OFC in favor of attributes in value space encoding in the area. LPFC and MPFC ROIs are outlined in orange and light blue respectively. All result are cluster corrected within the ROIs for FWE*<*0.05 with voxels surpassing p*<*0.001.

However, the extensive independent encoding of both attributes in value space and integrated value suggests these are separate neural computations. Part of this separability could originate from the great many raw attributes over which participant ratings (i.e., integrated value) might be computed, since we are necessarily only examining a small subset of such attributes. Thus, a linear combination over the attributes examined in the present study may not be sufficient to capture the full integrated value signal as it is represented in each individual. Accordingly, a difference between these representations may simply relate to this unexplained variance in integrated value arising from attributes outside the scope of our consideration.

## Discussion

This study investigated the neural mechanisms underlying the computation of integrated value ratings for multisattribute objects, a process central to flexible decision-making. We hypothesized the existence of “attributes in value space” as an intermediate computational step that bridges static stimulus-attributes and goal-dependent integrated values. While the first and final steps of this process have been the topic of much previous work (e.g. Kahnt et al. [2011], Iigaya et al. [2020]), the potential for the intermediate step necessary to flexibly transform a visual attribute to a object-based value judgment had not been directly explored. Through a combination of behavioral experiments and fMRI, our findings provide significant support for this hierarchical model of value construction, demonstrating distinct neural representations for raw attributes, attributes in value space, and integrated value ratings.

Our behavioral results clearly demonstrate the dynamic nature of integrated value. Participants significantly modulated their liking ratings for the same clothing items based on the instructed goal-context (ski trip, tropical island, or job interview). Successfully demonstrating a goal-dependent shift in valuation is necessary to distinguish between stimulus-intrinsic attributes and their value. Employing individualized attribute models, we derived estimates of attributes in value space through goal-specific weighting of raw attributes. Crucially, we found an important distinction between low-level attributes and high-level, subjectively-rated attributes wherein only high-level attributes generalized well to left-out data. This finding both speaks to the predictive power of high-level attributes in capturing general valuation, and provides further validation for a hierarchical model of value construction.

Turning to the fMRI data, we found evidence for a distributed architecture supporting hierarchical value processing. Raw attributes were robustly encoded in the visual cortex, consistent with their stimulus-intrinsic nature. The prefrontal cortex was hypothesized as a potential region to support the transformation of raw attributes into goal-dependent attributes in value space. Consistent with this hypothesis, we found that regions within both lateral and medial prefrontal cortex demonstrated a significant shift in the representation of attributes across different goal-contexts. This context-dependent neural coding, whereby a model trained on an attribute under one goal performed poorly when tested on another, precisely matches the expected behavior of attributes in value space—signals that are modulated by current goals.

We then sought to provide further neural evidence supporting the novel computational step we put forth as part of multi-attribute valuation. We directly investigated the presence of attribute-encoding in value space, alongside the existence of an integrated value signal within the prefrontal cortex. We found evidence for separable encoding of these different value representations. Integrated value signals were predominantly found along the medial wall of the prefrontal cortex (stretching towards the anterior cingulate cortex), while attributes in value space showed a more ventral representation along the medial wall and a dorsal-to-anterior pattern in lateral prefrontal cortex. We additionally found significant encoding of attributes in value space in regions including the medial and lateral orbitofrontal cortex (OFC). This distinct neural topography for attributes in value space and integrated value, however supports the possibility that these are two separable computational components in the flexible valuation process. As it was not our aim to be exhaustive in the set of attributes to decompose our stimuli into, this additional integrated value encoding might represent other attributes in value space not included in our set. Contrasting the neural representation of these attributes in value space with integrated value additionally opens up future research questions about the composition of integrated value. We hypothesize the potential of a more distributed representation of individual attributes in value space (where individual attributes in value space are encoded by partially overlapping but distinct sub-regions of the ventral prefrontal cortex), with this information combined into an integrated value signal for decision making and comparison. The fact that integrated value signals generalize well across goal-contexts further enforces its separable and abstracted nature, serving as a unified decision variable. While early theoretical work by von Winterfeldt and Fisher [1975] posited the potential for computing single attribute utility functions, direct neurobiological evidence for such value-like computation at the individual attribute level has remained elusive. Our findings, by demonstrating a neural correlate for attributes in value space, provide a critical missing piece. This work connects with previous research across various sub-domains of value computation; for instance, McNamee et al. [2013] reported a ventral-to-dorsal separation along the medial wall of the prefrontal cortex in their study of category-dependent value signals. In their work, participants’ subjective value for objects within specific categories (e.g., food, trinkets) was reflected in more ventral mPFC regions, while a more dorsal mPFC area held a category-independent value signal. Analogously, our study also reveals a gradient along the mPFC, where more ventral regions represent individual attribute values in a goal-dependent manner, and more dorsal regions encode the integrated value of overall objects. Both sets of results collectively underscore the brain’s capacity for hierarchical abstraction, moving from specific, context-dependent object properties to a more generalized, comparable value signal to facilitate efficient decision-making.

Our findings are broadly consistent with a recent study by Castegnetti et al. [2021], which observed that activity in the ventromedial prefrontal cortex (vmPFC) correlated with an item’s “usefulness” for the current goal, and that the neural code supporting this usefulness representation in the vmPFC was maintained and generalized across different goals. This finding resonates with our own report that integrated value signals, predominantly found along the medial wall of the prefrontal cortex, generalize well across goal-contexts, serving as a unified decision variable. In parallel, Castegnetti et al. [2021] also showed that while the dorsolateral prefrontal cortex (dlPFC) represented usefulness, its coding was specific to the current goal and did not generalize across contexts. The present study builds on those previous findings by providing a mechanistic account of how a context-specific value code can be produced in the first place through the transformation of attributes from raw stimulus features through to context-dependent attribute values which ultimately yields an integrated context-specific value representation. Collectively these findings shed light on the sophisticated interplay within the prefrontal cortex, where some sub-regions maintain generalized value signals, while others provide highly specific, context-modulated attribute representations, collectively underpinning the brain’s capacity to construct value on the fly and enable adaptive behavior in dynamically changing environments. It is important to acknowledge several limitations of the present work. First of all, while our experimental paradigm successfully manipulated goal-context, the scope of attributes and goals investigated was constrained. Other attributes will undoubtedly vary with our goal manipulation and many more will vary when considering other goals. However, given the overlap found across stimulus types for similar valuation processes, we expect our findings to generalize to other attributes, object classes and goals. Follow-up research could explore the specific intricacies that might exist under other conditions/objects/goals. Additionally, our research does not address how goal-dependent (attribute) values are formed over time. We might hypothesize that experience with items consisting of similar attributes under various conditions/contexts could inform current attribute valuation. Probing the mechanisms of such a learning process on the attribute-level could be an important direction of future research. Lastly, our findings on the relationship between attributes in value space and integrated value suggest a combination of overlapping and unique representations. Probing the extend of this overlap at various levels of neural computation (i.e., voxels to individual neurons) will require further research.

This study provides a comprehensive account of the neural computations underlying flexible value construction. By proposing and providing evidence in support of the existence of “attributes in value space”, we bridge the gap between static stimulus properties and dynamic, goal-dependent object valuation. Our findings highlight a hierarchical process, where sensory areas represent raw attributes, and prefrontal regions transform these into goal-sensitive attributes in value space, culminating in the computation of an integrated value signal that is robust and generalizable across contexts. This work advances our understanding of how the brain constructs flexible subjective value, a fundamental process for navigating a complex and ever-changing world.

## Materials and Methods

### fMRI experiment

Thirty-five participants (age 18 to 64 years, twenty female) participated in this experiment. Participants were recruited from the greater Pasadena (CA) area and were screened for MRI compatibility. Participants were paid $55 on average. All participants completed the study successfully and are included in the analyses.

#### MRI task procedure

Participants first answered a short demographics questionnaire after which they received the task instructions. These instructions included a more elaborate description of the goal-context to ensure a common understanding across participants. Following the task instructions, a set of ten practice trials was completed by each participant. During these practice trials participants were shown stimuli and a goal-context that were not used in the main experiment. This completed the pre-scanner portion of the experiment and participants were placed inside the scanner. When in the scanner participants were invited to browse through the instructions one more time after which the main portion of the experiment was started. Two sets of stimuli were created each consisting of various clothing items, one of which was female and the other male gendered. Stimuli are collected from the DeepFashion dataset and selected to not include any persons (see Liu et al. [2016]). Participants saw the set of stimuli that best matched their gender identity. In addition to these clothing stimuli a symbol and text label identifying the current goal-context were presented. Trials were presented in blocks of fifteen, within which the goal remains constant. Participants were shown the clothing item stimulus and a reminder of the goal, after which participants reported how much they would like to wear the clothing item with the current goal in mind (see Figure 1). Three blocks of fifteen images within a context were shown consecutively within an fMRI run. Each such fMRI run took 7 minutes and a total of five runs was completed by each participant (one participant completed an additional sixth run). A circular rating screen was used on which the rating 1 through 7 was displayed. The participants could navigate around this screen by pressing a left and right button to move either counterclockwise or clockwise respectively. Once they had reached the number corresponding to their intended rating they pressed a third button to confirm their rating. While the order of numbers and the starting location to navigate from was kept consistent the numbers were rotated around the screen randomly every trial. This setup helps dissociate between button pressing and the value rating inputted. For a traditional linear scale the ends of the scale (more button presses) are often associated with more extreme ratings. Additional correlation can arise since pressing left is often associated with the lower end of the scale while right is associated with higher ratings. Using the circular scale avoids these potentially confounding effects in the decoding analyses.

#### High-level attribute ratings

After completing the in-scanner task, participants were asked to give high-level attribute ratings for the stimuli used. This rating was self-paced and took around 20 minutes to complete. The high-level attributes rated were ‘warmth’, ‘formalness’, ‘festiveness’, ‘youthfullness’ and ‘comfort’ (youthfullness attribute was excluded from analyses due to correlation with other attributes - see supplement Figure S2F).Participants rated each stimulus they saw during the experiment on five high-level attributes on a rating scale from 1 to 10. One subject had no variance in their provided comfort ratings across stimuli, this subject is therefore excluded from any comfort-attribute based models.

### Online experiment

The online sample consisted of 95 participants (57 female) and was collected through Prolific. Online participants were paid approximately $5 on average.

#### Online task procedure

Participants performed a rating task under four instructed goal-contexts (tropical island vacation, ski trip, job interview and summer wedding). Participants saw 100 unique, gender-matched stimuli total, resulting in 400 total trials (see Figure S1 for online design). Participants indicated how much they would like to wear the item shown given the goal-context on a scale from 1 through 9 (1; don’t want to purchase at all, 9; want to purchase a lot). This rating was reported using the 1 through 9 keys at the top of a participants keyboard. Image order was fully randomized and trials under a certain goal-context were presented in a blocked manner (20 trials before switching). Trials were self-paced and participants moved on to the next stimulus-context combination when a rating was entered. The fourth goal condition included in the online experiment was not in the fMRI study due to time constraints in the scanner. This goal-condition is therefore excluded from further analyses.

### Behavioral analyses

Note that integrated value and raw attribute ratings are independent of gender therefore data from both genders was analyzed jointly (see Supplemental Figure S2B, D&E).

#### Low-level attribute extraction

Low-level computer vision attributes were extracted using the same methods as those used in Iigaya et al. [2021]. These attributes capture both segment-specific and overall stimulus characteristics. Please see Iigaya et al. [2021] and Li and Chen [2009] for further details.

#### Mixed-effects modeling: attribute main effects

For each attribute in each context independently we ran linear model predicting value ratings. This model included the attribute and a random intercept for each subject (value = attribute + (1—subjectID)). We then evaluated the significance of the attribute coefficient. This set up resulted in 4 attributes x 3 contexts models.

#### Beta weight estimation modeling

High-level raw attribute beta weights (shown in Figure 2B&C) were fitted within subject for each context separately. The model used in this analysis included the four high-level attributes and was predicting the object value ratings of the participant and context in question. Since the online participants did not provide subjective attribute ratings, we instead used the stimulus’ mean attribute rating provided by the MRI participants. Models were fit separately within subject and context to ensure individual and within goal differences were maximally captured. This is especially important since we would expect such inter-individual differences to also be reflected on a neural level. For example, for a participant whose value rating is highly dependent on the comfort of the item we would expect a stronger comfort representation than for a participant whose value ratings are independent of comfort.

Since the distribution of value ratings displayed a bimodal pattern we additionally repeated this analysis using a logistic regression model on binarized (median split) value ratings. The beta weights fit using this approach are shown to correlate strongly with those fit using the linear model (see Supplemental Figure S2G).

#### Leave-one-out behavioral generalization modeling

We used an elastic net model (alpha=0.2 and ratio=0.5) trained to predict value ratings on all but 1 left-out participant. The model was then used to predict value ratings for the left-out participant. The correlation between these predicted ratings and the true ratings was used to evaluate the model. This procedure was repeated for each participant in the sample. Similarly, we trained an elastic net model to all participants within one gender-group and tested on one of the opposite-gender participants. This process was repeated until each subject had been left out once.

### Neural analyses

#### Data preprocessing

fMRI data were preprocessed using a custom pipeline in the Nipype (Gorgolewski et al. [2011]) environment. Functional images were extracted using functions from the Nilearn (Abraham et al. [2014]) toolbox and motion corrected using the MCFLIRT algorithm (Jenkinson et al. [2002]) from the FM-RIB Software Library (FSL; Jenkinson et al. [2012]), which estimated six motion parameters to be included as covariates in the statistical model along with estimated motion outlier volumes. Other covariates were estimated using a principle components analysis on both the timeseries of white matter and CSF voxels, demarcated with the co-registered T1 structural image, as well as outlier timeseries using the CompCor algorithm (Behzadi et al. [2007]). The functional data were unwarped to correct for distortion using FSL’s TOPUP algorithm (Andersson et al. [2003]) with reverse phase-encoded spin-echo EPI images. Both T1 and T2 structural images were corrected for field bias using the N4 algorithm (Tustison et al. [2010]) and brain extracted using Advanced Normalization Tools (ANTs; Avants et al.). Affine co-registration transformations were estimated between the functional image and the T2 as well as between the T2 and T1 structural images. A non-linear transformation was estimated to normalize the T1 structural to the MNI152 template (Mazziotta et al. [2001]) using the symmetric normalization algorithm (Avants et al. [2008]). All transformation matrices were estimated using ANTs, and applied to the functional data simultaneously to normalize to template space. Finally, the functional data were de-spiked and smoothed with a 6mm FWHM kernel (smoothing was applied for encoding analyses only).

#### Univariate encoding of raw attributes

fMRI univariate encoding analyses were performed using FSL (Jenkinson et al. [2012]). Custom onset files created regressors spanning the stimulus-on period. The raw attribute analysis included the three contexts as regressors in addition to the four high-level attribute regressors and four low-level attributes (average saturation of the 2nd segment, average brightness of the 2nd segment, mean saturation of the 2nd segment and entropy of the largest segment). We then constructed two independent F-tests over the four high-level and the four low-level attributes. F-test statistics were computed based on the group-level results where voxels were thresholded at a z-score of 3.2 (*p <*= 0.001) and cluster correction was performed (*p <*= 0.05). The proportions of significant voxels across ventral and dorsal visual ROIs were computed as those passing the *p <* 0.01 threshold before cluster correction. ROIs were defined using the atlas from Wang et al. [2015].

#### ROI generalization decoding

Single-trial *ω*’s were extracted for each voxel from the un-smoothed neural data time-locked to the stimulus on period. These were then masked based on the ROI of interest. Visual ROI masks were created using Wang et al. [2015] atlas. Frontal ROI masks were created by merging regions specified in Tzourio-Mazoyer et al. [2020]. More specifically, the lOFC ROI was created by merging the frontal mid orbital ROIs, frontal inferior orbital ROIs and the frontal superior orbital ROIs. The mOFC consisted of the frontal medial orbital ROIs and the rectus ROIs. The dmPFC ROI consisted of the frontal superior medial ROIs ad the cingulum anterior ROIs. The dlPFC ROI was made up out of the frontal mid ROIs and the frontal superior ROIs. Lastly, the vlPFC consisted of the frontal inferior operculum ROIs and the frontal inferior triangularis ROIs. The larger LPFC and MPFC ROIs were created by merging the lOFC, dlPFC and vlPFC or mOFC and dmPFC respectively.

The generalization decoding was ran over each context combination, with each context considered the training context and each context considered as the test context (leading to a total of nine train-test combinations per variable of interest per ROI). Trials were split into training and testing sets using a 5-fold cross-validation scheme. Importantly, for the out-of-context decoding (training context is not equal to testing context) trials belonging to the training context were still subset using this approach to ensure equal training data for across and within context modeling. The decoding was performed using a ridge classification model. The dependent variable was median split to allow for binary classification. The average accuracy across folds was returned for model evaluation.

Test set group-level accuracy means were compared to permuted chance distributions. Chance distributions were computed through shuffling the dependent variable, retraining the classifier model and evaluating its performance using the same metrics as non-permuted. We computed 500 such shuffled model accuracies (across 5-folds each) for each train-test context pairing for each ROI and each attribute. Between context statistics were computed using T-tests for the differences for each train-test pair shown.

Additionally we computed subject-level decoding accuracy means across all train-test combinations for each ROI and attribute. We split these mean accuracies into those for within-context models and out-of-context models. The significance of the difference between within vs out-of-context models was computed using Wilcoxon signed rank tests.

#### ROI decoding of integrated valu

ROI-based decoding analyses were used to determine the presence of multi-variate integrated value representations across the whole task for the ROIs introduced above. For this analysis we again binarized the dependent variable (median split). We perform stratified 3-fold cross validation and trained a ridge classifier model for each fold. The mean accuracy across the cross-validation folds was saved for each ROI for each participant. Significance of the group-level mean was again compared to permuted null distribution. We again computed 500 permuted model accuracies for each individual (and 3-folds cross-validation within each shuffle). We then computed group level permuted accuracies for comparison.

#### Value-based encoding models

Univariate integrated and attribute value regression models were set up as similarly as possible. All these models included three context regressors (ski, job and island context). In addition, the models of integrated value included a parametrically modulated regressor capturing either integrated value ratings reported by the participants (or behavioral model estimates for the integrated value - see supplemental Figure S7).

For the attribute in value space model we use the same three context regressors in addition to 4 attribute in value space parametric regressors. These attribute in value space regressors were created by taking the subjective attribute ratings multiplied with subject specific *ω* weights from the current-context’s behavioral GLM. For example, if the trial stimulus is a jacket in the job interview context the *ω* weights belonging to the attribute regressors in the job interview context GLM is used. Since the attributes in value space model included 4 regressors of in interest (one for each of our a priori high-level attributes) we additionally computed a contrast taking the mean across individual regressor effects. Similarly to the raw attribute encoding models, we thresholded individual voxels at *p <* 0.001 and cluster corrected to *p <* 0.05.

For the models including both integrated value and attributes in value space we combined the following regressors together: integrated value ratings, 4 attribute in value space regressors and three context regressors. The integrated value and the 4 attribute in value space regressors might be modeling overlapping variance. In extremes, two perfectly correlated constructs in one analysis should result in a null result when testing contrasts to identify one or other type of representation. As due to the extra sum of squares principle (Draper and Smith [1998]), all shared variance (i.e. which in such a case would be all of the variance) between these representations should have been partialed out.

