## Supplemental information for "The neural computations underlying context dependent attribute-based valuation of complex stimuli"

### Supplementary Information

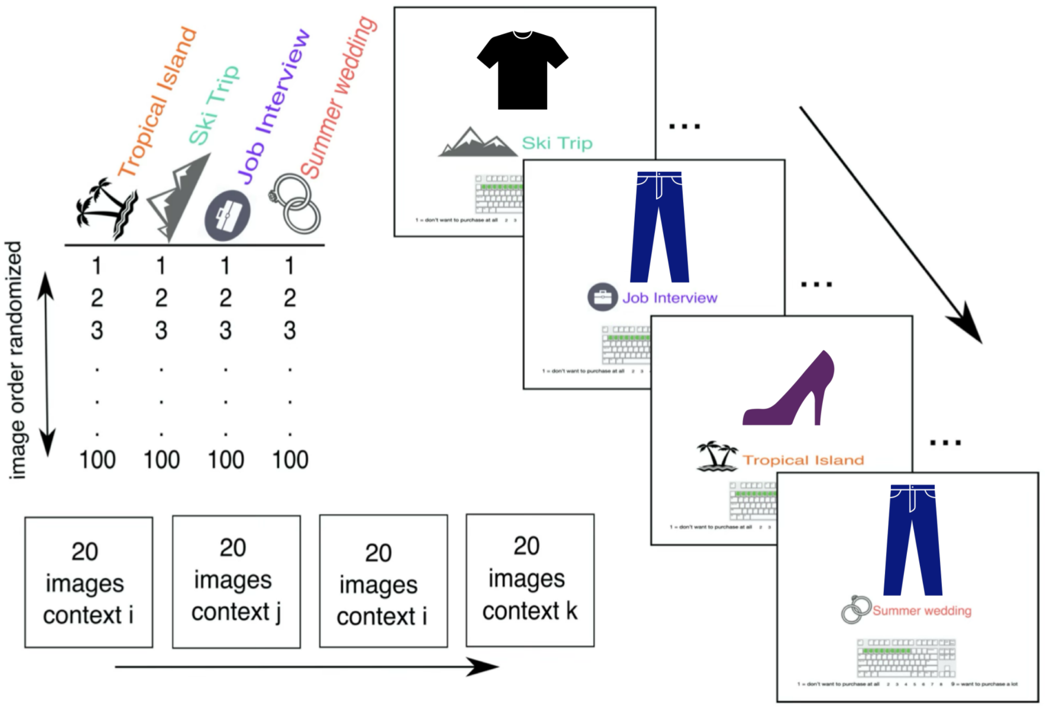


**Figure S1:** Online experiment design. Participants performed a rating task under four instructed goal-contexts (tropical island vacation, ski trip, job interview and summer wedding). Participants indicated how much they would like to wear the item shown given the goal-context on a scale from 1 through 9 (1; don't want to purchase at all, 9; want to purchase a lot). This rating was reported using the 1 through 9 keys at the top of a participants keyboard. Image order was fully randomized and trials under a certain goal-context were presented in a blocked manner (20 trials before switching). Trials were self-paced and participants moved on to the next stimulus-context combination when a rating was entered. Images shown are placeholders, actual stimuli were from Liu et al. [2016].

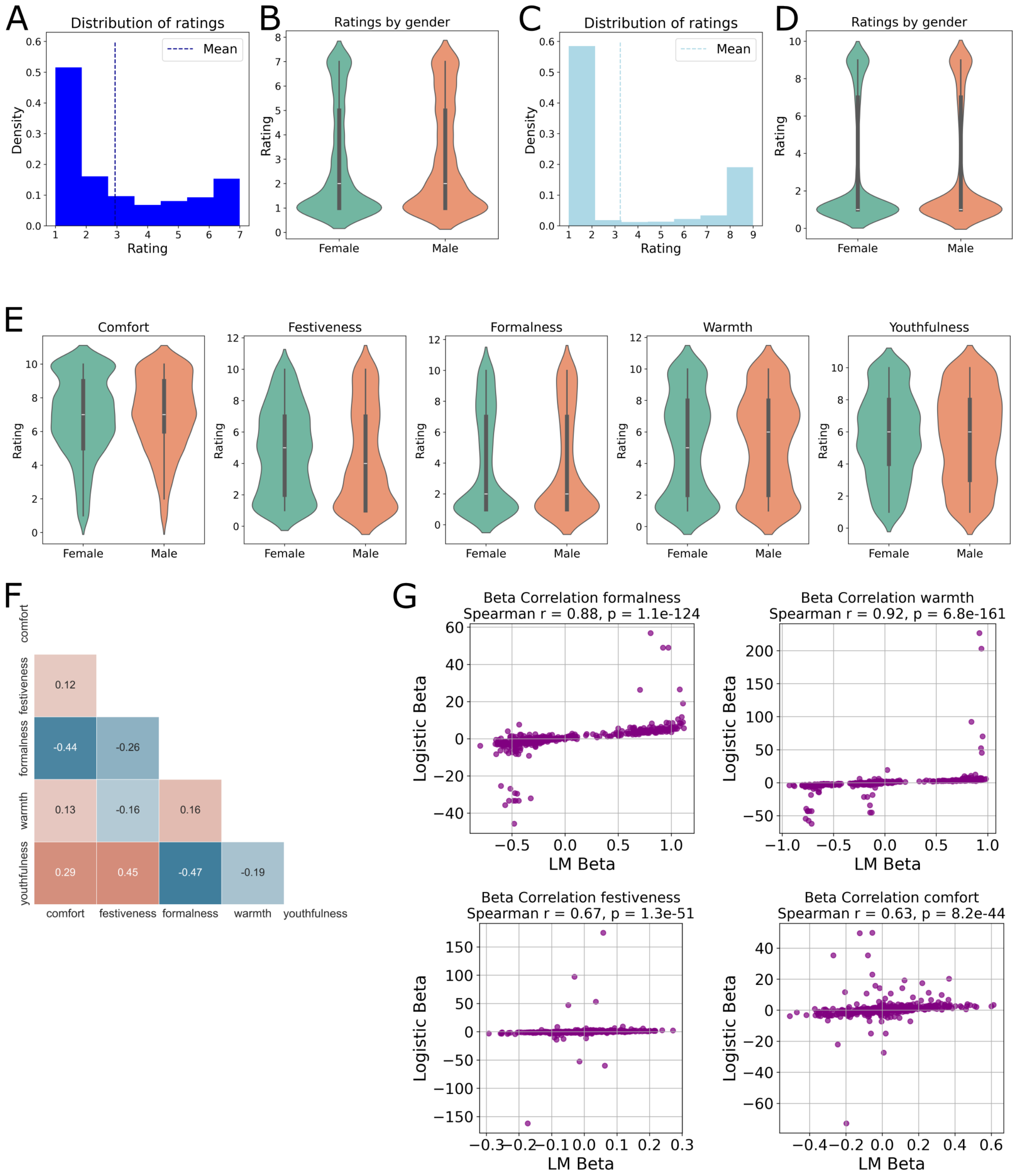


**Figure S2:** **A)** Distribution of ratings across participants and contexts for the MRI sample (n=35). **B)** Ratings split by gender for the MRI sample. **C)**Ratings distribution for the online sample. **D)** Ratings split by gender for the online sample. **E)** Attribute ratings split by gender. Note this includes only the MRI sample since no attribute ratings were obtained for the online sample. **F)** Average pair-wise correlations for all attributes rated by the MRI sample. **G)** Spearman rank correlation. Outliers (z¿3) removed for plotting only.

| Context | Attribute | Estimate | Standard error | T-value | P-value |
|---|---|---|---|---|---|
| Island | warmth | -0.398 | 0.011 | -36.4 | 1.115605e-234 |
| Island | comfort | 0.167 | 0.019 | 8.7 | 4.082655e-18 |
| Island | formalness | -0.236 | 0.012 | -19.7 | 1.065373e-80 |
| Island | festiveness | 0.146 | 0.016 | 9.2 | 5.781847e-20 |
| Ski | warmth | 0.486 | 0.010 | 48.1 | 0.000000e+00 |
| Ski | comfort | 0.3119 | 0.019 | 16.5 | 4.469856e-58 |
| Ski | formalness | -0.098 | 0.013 | -7.5 | 1.092899e-13 |
| Ski | festiveness | -0.057 | 0.016 | -3.4 | 5.804354e-04 |
| Job | warmth | 0.090 | 0.014 | 6.4 | 1.559250e-10 |
| Job | comfort | -0.380 | 0.019 | -20.2 | 4.266328e-84 |
| Job | formalness | 0.561 | 0.008 | 72.9 | 0.000000e+00 |
| Job | festiveness | -0.209 | 0.016 | -12.8 | 2.047208e-36 |

**Table S1:** Attribute main effects across subjects within each goal-context.

| Raw attribute | Kruskal - Wallis H | P-value | Group |
|---|---|---|---|
| Formalness | 70.7 | 4.45E-16 | fMRI |
| Warmth | 84.9 | 3.72E-19 | fMRI |
| Festiveness | 4.6 | 0.10 | fMRI |
| Comfort | 15.6 | 0.0004 | fMRI |
| Formalness | 176.70 | 4.35E-39 | Online |
| Warmth | 212.0 | 9.18E-47 | Online |
| Festiveness | 6.8 | 0.03 | Online |
| Comfort | 8.1 | 0.02 | Online |

**Table S2:** Statistics of Kruskal - Wallis tests for coefficients fit using the Logistic model.

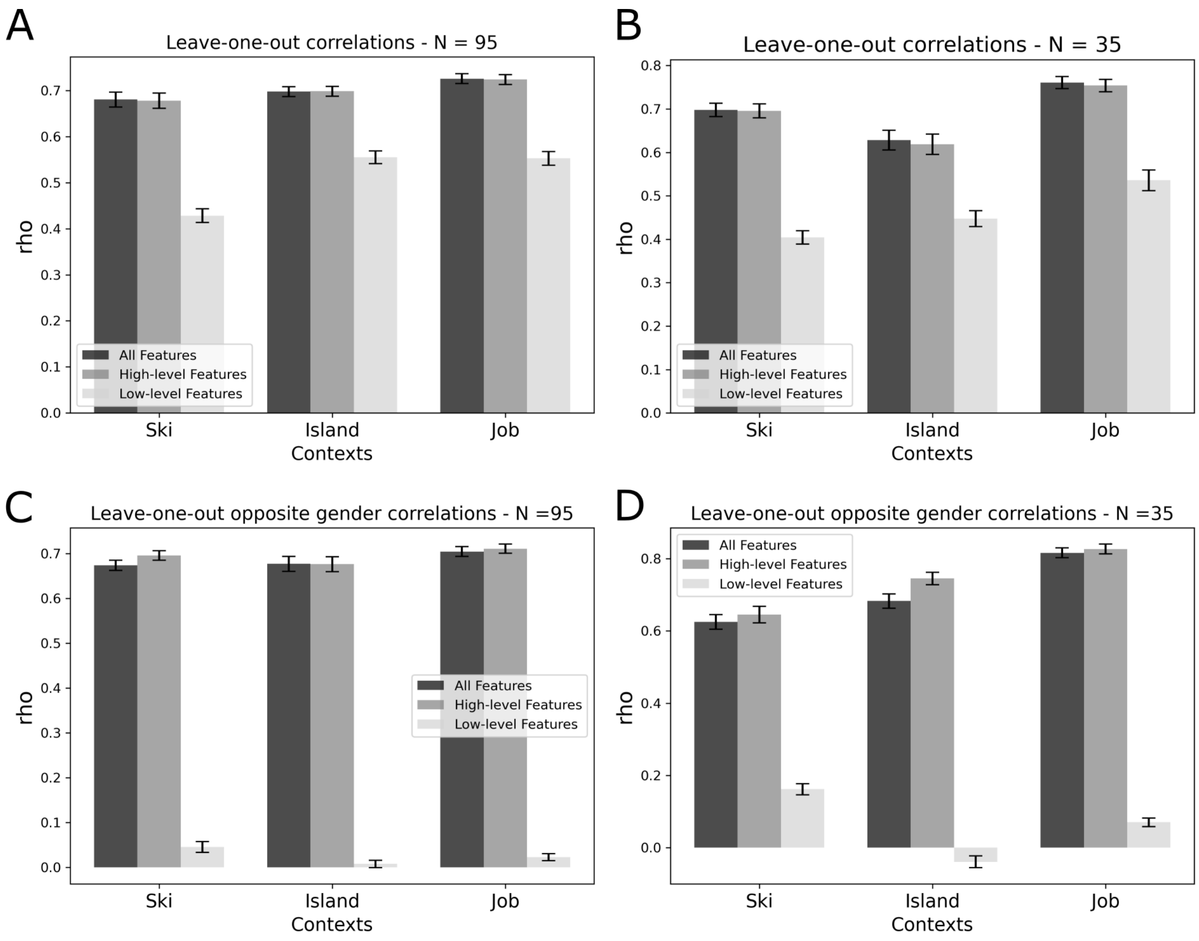


**Figure S3:** **A)** Leave-one-out model fits as in Figure 2D but for only the Online experiment. Generalization across individuals, with models trained on N-1 and tested on left out 1 subject. **B)** Across subject generalization performance for fMRI subjects. **C)** Leave-one-out generalization results as in Figure 2E. Model trained on all subjects from one gender and tested on each of the opposite gendered subjects one-by-one. Results shown for Online experiment only. **D)** Across-gender generalization results for fMRI experiment group.

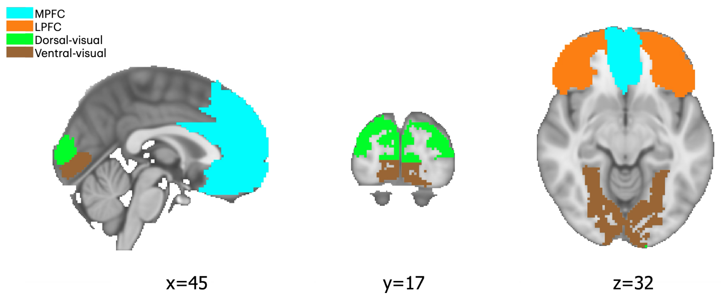


**Figure S4:** ROI locations as used in generalization analysis. The two visual ROIs are derived from Wang et al. [2015] and are shown in green and brown. Frontal ROIs are from Tzourio-Mazoyer et al. [2020] shown in blue and orange. For more details see Methods.

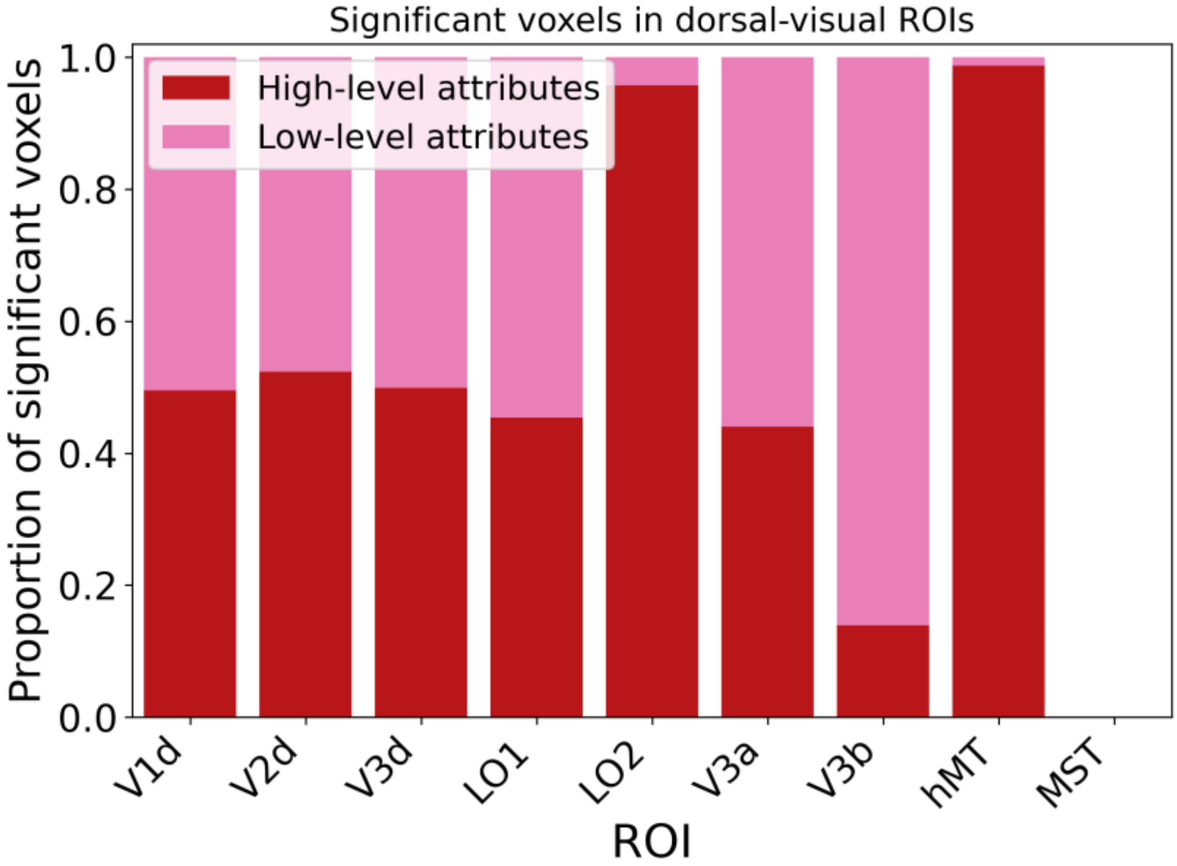


**Figure S5:** Proportion of significant voxels along posterior-to-anterior ROIs in the dorsal visual stream.

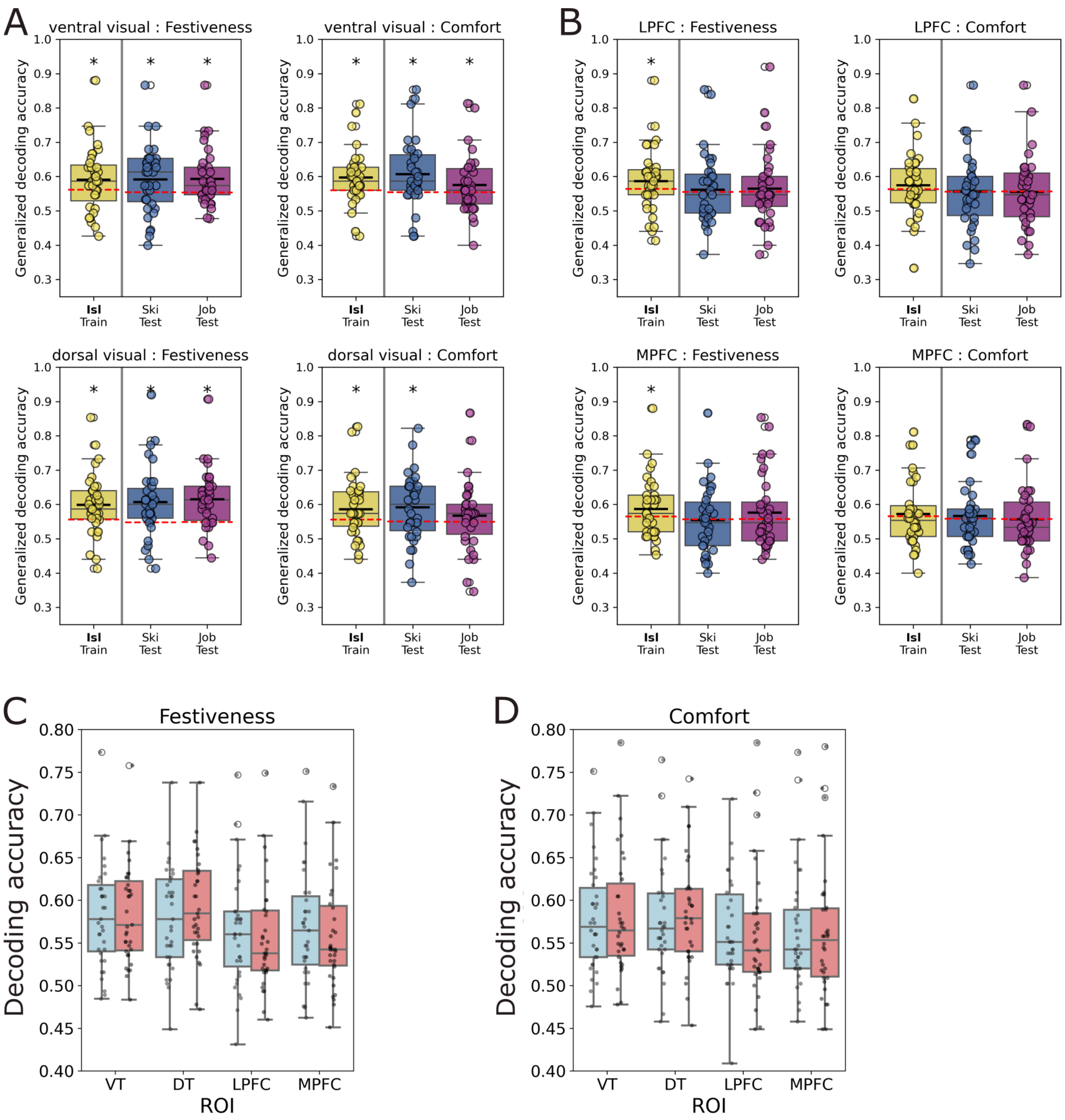


**Figure S6:** **A)** Generalization decoding accuracies for festiveness and comfort in ventral and dorsal visual ROIs. We observe good generalization across attributes and ROIs. **B)** Generalization accuracies for frontal ROIs for festiveness and comfort. We see only festiveness can be decoded in the training context. Additionally, we observe no significant generalization across ROIs for both festiveness and comfort. **C)** Averaged decoding accuracies for all train-test combinations show a similar pattern as **A**. We observe no differences between in- and out-of-context decoding accuracies. **D)** Similarly for the comfort attribute, we observe no differences in decoding accuracies between models tested on the training context and those tested on another context.

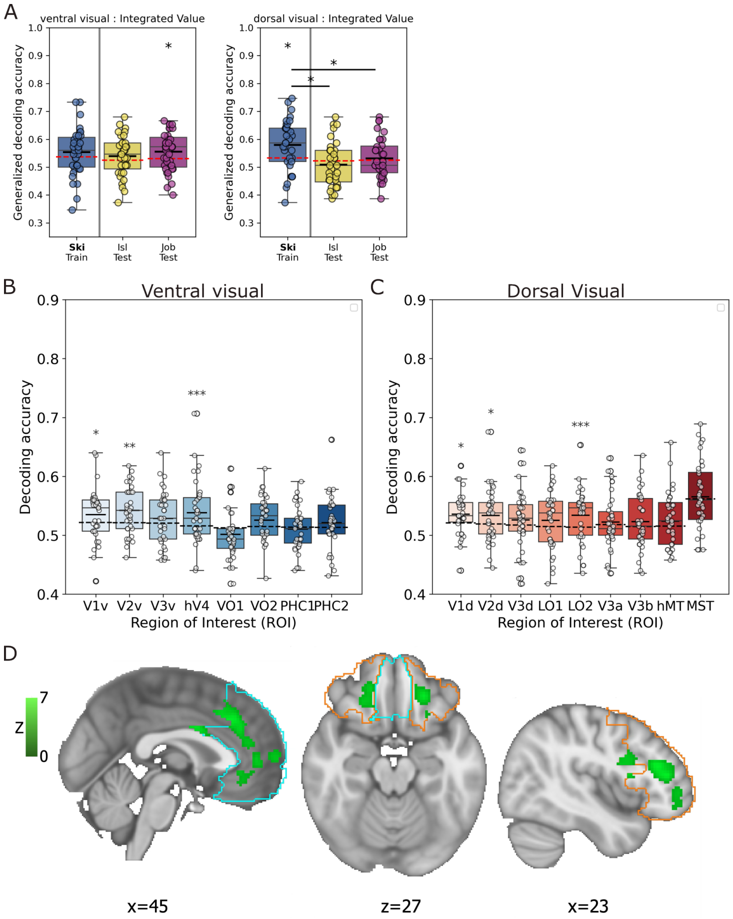


**Figure S7: A)** Generalization decoding for integrated value for ventral and dorsal visual ROIs. We observe no significant decoding in the training context for the ventral visual ROI, indicating there is no information about integrated value held in this region. Additionally, while we do observe significant decoding in the training context for the dorsal visual ROI, this decoding does not generalize the the test contexts. **B)** Within subject decoding accuracies of integrated value for each of the ROIs along the ventral visual pathway. For these decoding analyses all trials across goal-contexts were used (and train-test splits were set up, see Methods). Box-and-whiskers show across subject medians and quartiles similar to previous figures. Significance of these group-level accuracies is computed through comparison to a permuted null and indicated as follows: $*$: $p<0.05$, $**$: $p<0.01$, $***$: $p<0.001$. **C)** Similarly, for the ROIs along the dorsal visual pathway we show the within-subject decoding accuracies and group-level statistics for the integrated value ratings across the whole task. **D)** Encoding model of integrated value using the behavioral model estimates from the linear model (see Methods) instead of the participant rating. Max voxel in LPFC: [25, 81, 43], z=6.5
